# Exploring Volatile Sulfur Production by Arctic Soil Bacteria and Emissions from Thawing Permafrost

**DOI:** 10.64898/2026.05.23.727374

**Authors:** Kajsa Roslund, Miguel Salinas-García, Anders Priemé, Riikka Rinnan

## Abstract

Warming of the Arctic enhances microbial activity and the decomposition of large stocks of organic matter retained in permafrost soil. Resulting changes in the availability of sulfur may lead to increased emissions of volatile sulfur compounds (VSCs), which impact atmospheric particle and cloud formation, terrestrial and aquatic acidification, and malodor. Marine microbial production of dimethyl sulfide (DMS) has been studied for decades but other VSCs have been largely ignored, while VSC emissions from terrestrial ecosystems are even less studied. Currently, we lack fundamental understanding of the metabolic processes behind VSC production in permafrost soil bacteria, essential for estimating how emissions may change due to thawing. To fill this knowledge gap, we measured VSC emissions from thawing permafrost and three bacterial strains isolated from Greenlandic permafrost and biological soil crust. We show that the bacterial strains produced high levels of VSCs *in vitro* – including hydrogen sulfide, methanethiol, DMS, dimethyl disulfide, and dimethyl trisulfide. We further show that the same VSCs were also emitted from permafrost upon thaw. Metabolic pathway mapping of the bacterial strains revealed both inorganic sulfate reduction pathways and amino acid metabolism behind bacterial VSC production. High production of VSCs in the late-active and stationary phase suggests connection to secondary metabolism, except for DMS which was linked to early growth, and possibly, primary energy metabolism. Our findings suggest that thawing increases VSC emissions from permafrost soil, possibly leading to higher input of sulfur into the atmosphere from the warming Arctic in the future.

## 1. Introduction

Permafrost covers approximately 15% of the land surface in the Northern Hemisphere (Obu, 2021), and soils in the permafrost region store an estimated 1300–1700 Gt of organic carbon – more than the current amount in the atmosphere (Hugelius et al., 2014; Schuur et al., 2015). Organic matter in permafrost soil remains largely undecomposed because the frozen conditions limit microbial activity. However, the Arctic warms three to four times faster than the global average (Rantanen et al., 2022), causing accelerated permafrost thaw and a more favorable environment for microbes. This may accelerate the cycling of greenhouse gases and volatile organic compounds (VOCs) (Miner et al., 2022), not only in permafrost soil, but also in biological soil crust – the uppermost layer of soil inhabiting complex communities of microorganisms and bryophytes. While the majority of the decomposable carbon revealed by permafrost thaw is released as carbon dioxide (CO_2_) and methane (CH_4_) (Nadeem et al., 2024; Turetsky et al., 2020), there is also increasing evidence of the release of VOCs from thawing permafrost (Kramshøj et al., 2018; Kramshøj et al., 2019; Li et al., 2020) and Arctic surface soil (Jiao et al., 2023; Jiao et al., 2025). However, little is known about cycling of volatile sulfur compounds (VSCs).

Human activity – including fossil fuel combustion, pulp and paper mills, wastewater treatment, and landfills – releases sulfur into the atmosphere mainly as sulfur dioxide (SO_2_), while microbial degradation of organic matter and volcanic emissions release mostly reduced VSCs (Holloway & Wayne, 2010). These include hydrogen sulfide (H_2_S), methanethiol (MeSH), dimethyl sulfide (DMS), and dimethyl disulfide (DMDS). VSCs are rapidly oxidized in the atmosphere to SO_2_ and ultimately converted into sulfuric acid (Berndt et al., 2023; Yu et al., 2024). As such, they impact the acidification of terrestrial and aquatic ecosystems (Sase, 2023) and take part in cloud formation, increasing the cloud albedo of our planet (Park et al., 2021). Currently, only a small portion of VSCs reaches the atmosphere as the majority is recycled in the soils and sediments where they are produced – carbon is converted to CO_2_ and biomass, while sulfur is retained as sulfate (SO_4_^2^), elemental sulfur, or metal sulfides (Lomans et al., 2002; Zhou et al., 2025).

The most studied aquatic and terrestrial VSC is DMS. The interest in DMS has been driven by early observations of elevated DMS concentrations in marine environments, produced by algae and phytoplankton, as well as its high atmospheric reactivity (Wohl et al., 2024). DMS emissions have also been observed from freshwater (Steinke et al., 2018), as well as terrestrial sources, such as drought-affected tropical soil (Pugliese et al., 2023). A recent study shows DMS emissions, alongside H_2_S and MeSH, from waterlogged, minerotrophic peatland (Lehnert et al., 2024). While DMS is arguably the most abundant VSC in the atmosphere, the importance of non-DMS VSCs is gradually being recognized, and in the marine environment, they can contribute 37% of the total gas-phase sulfur (Kilgour et al., 2022).

Bacteria produce VSCs mainly via reduction and methylation of inorganic sulfur or degradation of sulfur-containing amino acids and other organic sulfur compounds. SO_4_^2^ is reduced to H_2_S, which can be further methylated to organic sulfur compounds, such as DMS or MeSH (Li et al., 2023; Zhang et al., 2024). The genes encoding sulfate-reducing enzymes in bacteria are highly conserved and have existed for billions of years (Neukirchen et al., 2023), and today are commonly found in nature. In aquatic environments, the organosulfur compound 3-dimethylsulfoniopropionate (DMSP) is produced from organic matter mainly by algae, but also by some bacteria (Williams et al., 2019). DMSP is then cleaved to DMS and smaller amounts of MeSH by bacteria (Liu et al., 2022; Shao et al., 2019). Contrary to sulfate reduction and DMSP catabolism, bacterial degradation of sulfur-containing amino acids – methionine and cysteine – leads to a variety of VSCs, including H_2_S, MeSH, DMS, DMDS, dimethyl trisulfide (DMTS), and several thioesters (Schulz & Dickschat, 2007). Consequently, the composition of VSCs produced in an ecological niche reflects the type of metabolic pathways present in the microbial community, as well as the form of sulfur available in the environment.

Sulfate-reducing bacteria are integral part of microbial communities in hypersaline lake sediments (Foti et al., 2007), wetlands and soil under repeated flooding-drying cycles (Demin et al., 2024), as well as in thawing permafrost (Li et al., 2024; Stackhouse et al., 2015). However, beyond this, we know nothing about sulfur processing in bacteria inhabiting Arctic soils and permafrost. All the VSC production pathways could coexist depending on the structure of the microbial community, nutrient availability, and prevailing conditions, such as redox potential, temperature, and salinity. A shift in any of these factors may impact the availability of sulfur in the environment and the dominating metabolic pathways, which in turn determines which VSCs and how much is ultimately produced. The current lack of knowledge makes it impossible to estimate how VSC emissions will be impacted when temperatures continue rising.

We have aimed to fill this knowledge gap by investigated the VSC production capacities and metabolic pathways of three bacterial strains isolated from Greenlandic soil and permafrost. Concentrations of VSCs produced by mono- and cocultures were monitored throughout the bacterial growth to link the VSCs produced to different phases of the bacterial life cycle when sulfur-containing nutrients are available. Genomic data was used for metabolic pathway mapping to connect the VSCs observed to bacterial sulfur-cycling potential. Additionally, we analyzed VSC emissions from permafrost soil samples during thaw to examine if volatile sulfur is also released from naturally oligotrophic samples. We had four central hypotheses: (1) The studied bacterial strains produce reduced VSCs when sulfur is readily available in the environment, (2) the strains differ in both the composition and concentrations of VSCs in ways that can be explained by their sulfur metabolic pathways, (3) VSC production is connected to the bacterial life-cycle determined by activation of primary versus secondary metabolism, (4) thawing permafrost emits reduced VSCs originating at least in part from microbial production.

## 2. Materials and Methods

### 2.1 Bacterial cultures

Bacterial strains studied in this work – *Nesterenkonia aurantiaca* CMS1.6, *Oceanobacillus* sp. CF4.6, and *Arthrobacter arnarulunnguaqae* KK5.5 (Salinas-García & Priemé, 2026) – were isolated from soil (Cap Morris Jessup), biological soil crust (Citronen Fjord), and permafrost (Kap København), respectively, sampled from Northern Greenland. Sampling, isolation, and characterization of the strains have been described in detail elsewhere and their genomes are publicly available (Salinas García & Priemé, 2025). These strains were chosen based on our prior knowledge of them as producers of volatile metabolites (oma paperi). The chosen strains also represent traits relevant for Arctic regions, such as adaptation to extreme cold and high salinity (Dai et al., 2022; Shen et al., 2021; Yukimura et al., 2009) (Yukimura et al. 2009)(Doytchinov and Dimov 2022). Culturing was done in a modified HM medium, which was designed for moderately halophilic bacteria (Ventosa et al., 1982). The full composition of the medium can be found in Supplementary Material (**Table S1ab**), with sulfur available as amino acids and sulfate to promote VSC production.

Culture protocol is described in detail in Supplementary Material. Briefly, isolates were streaked onto agar plates until colonies formed. One colony was transferred to liquid media (preculture) and incubated overnight. Samples for VSC analysis were prepared by transferring 600 µL of the precultures (200 µL of each for the coculture) into 20 mL of sterile media in a 330 mL glass jar with a double-inlet cap. Triplicate samples (N=3) were prepared for each strain, coculture, and control (sterile nutrient media). Growth curves (**Fig. S1a**) were measured from parallel cultures to avoid contamination in the VSC measurements. Relative amounts of strains in the coculture were determined via serial dilution and colony counting (**Table S2**).

### 2.2. Permafrost samples

Sampling, processing, and characteristics of the permafrost soil as well as their VOC emissions other than sulfur-containing compounds have been described in detail previously (Kramshøj et al., 2018). Briefly, nine independent permafrost cores were collected in 2015 in Western Greenland, 10 cm below the permafrost table. The samples were homogenized and stored frozen for approximately six months before analysis. Six of the nine replicate samples of frozen permafrost, 13 g each by fresh weight, were placed in 200 mL glass jars with double-inlet caps. An additional empty jar was used as a control.

### 2.3 Analysis of VSCs

#### 2.3.1 Continuous headspace measurements

The measurement setup for culture headspace measurements has been described in detail previously (Salinas-García et al., 2025). Briefly, bacterial culture headspace was sampled continuously with a flow rate of 70 mL min^-1^, using VOC-free air as the carrier gas, at 25 °C and with agitation. A commercial proton-transfer-reaction mass spectrometry (PTR-TOF-MS) instrument (PTR-TOF 1000 ultra, Ionicon Analytik, Austria) was used for real-time monitoring of VSC concentrations in the culture headspace, with hydronium (H_3_O^+^) as the reagent ion and measurements performed from the mass-to-charge ratio (*m/z*) 17 to 239 with a 0.25 Hz frequency. An automated valve-system was used for switching between the bacterial cultures throughout the culturing period (68 h). Operational parameters are specified in Supplementary Material.

The measurement setup for permafrost headspace measurements has also been described in detail before (Kramshøj et al., 2019). Briefly, permafrost headspace was sampled continuously with a flow rate of 100 mL min^-1^, using VOC-free air as the carrier gas, at 6 °C. Another commercial PTR-TOF-MS instrument (PTR-TOF 8000, Ionicon Analytik, Austria) was used for measuring VSCs from thawing permafrost with H_3_O^+^ as the reagent ion, and measurements performed up to *m/z* 280 with a 1 Hz measurement frequency. Operational parameters have been specified previously (Kramshøj et al., 2019). An automated valve-system was used for switching between the permafrost samples throughout the thawing period (40 h).

Raw spectra were processed with the PTR-MS Viewer program v. 3.4.4 (Ionicon Analytik, Austria). The PTR-TOF-MS allows for tentative identification of peaks by comparison of the measured accurate mass to the theoretical exact mass. In this study, we identified five VSCs based on their accurate masses (**Table S3**), with the identity accepted if the mass difference (*Δm*) was <0.010. Further confirmation was done via isotopic pattern analysis (PTR-MS Viewer program v. 3.4.4, Ionicon Analytik, Austria). Peak ion counts-per-second (cps) were converted with the PTR-MS Viewer program into volume mixing ratios (c_VMR_), referred to here as concentrations, in parts-per-trillion (ppt, 10^-12^ = pmol mol^-1^).

An average concentration of each VSC at each time point during the culture period was calculated as the mean ± standard error (SE) of the triplicate samples. Blank (sterile nutrient) concentrations were subtracted from bacterial samples. Emission rate (pmol h^-1^ mL^-1^) at each time point was calculated by multiplying the concentration (ppt) by the molar flow rate through the culture headspace (mol h^-1^) and dividing by the culture volume (mL). For growth normalized emission rates (pmol h^-1^ 10^-8^ cells), values were normalized to OD_600_ = 1·10^8^ cells mL^-1^. Total emissions (pmol mL^-1^) were calculated by integrating emission rates over the whole 68 h culturing period.

For the permafrost samples, an average concentration of each VSC at each time point during the thawing period was calculated as the mean ± SE of the six replicates. Blank (empty jar) concentrations were subtracted from permafrost samples. For emission rates (pmol h^-1^ g^-1^), VSC concentrations (ppt) were multiplied by the molar flow rate through the jar headspace (mol h^-1^) and divided by the dry weight of the permafrost sample. Total emissions (pmol g^-1^) were calculated by integrating emission rates over the whole 40 h thawing period.

#### 2.3.2 Gas chromatography–mass spectrometry (GC–MS) identification of VSCs

The headspace gas from each bacterial strain and coculture was sampled once during the culturing (at 22-54 h depending on the strain) into sorbent tubes (Tenax TA + Carbograph 1TD, Markes International, UK) for 5 min with a flow rate of 200 mL min^-1^. The sorbent tubes were stored at 4 °C with storage caps on until analysis to avoid loss of analytes. The sorbent tubes were thermally desorbed (TD100-xr, Markes International, UK) and VSCs analyzed with GC-MS (7890A + 5975C MSD, Agilent Technologies, US) with helium as carrier gas. Operational parameters are specified in Supplementary Material.

Identities of three of the heavier VSCs (DMS, DMDS, and DMTS) were confirmed by comparing the measured spectra to standard compounds and to a spectral library (NIST23, Gaithersburg, US). The identity was accepted if the retention time was within ±0.05 min compared to the standard compound and the match factor and the reverse match factor between the measured and reference spectra were >700 and the probability of the identity >30% (**Table S4**). MeSH and H_2_S are too volatile to be captured with the sorbent tubes used in this study and quantification and identification was performed with PTR-TOF-MS as described above. Permafrost samples were not analyzed with TD-GC-MS.

### 2.4 Analysis of sulfur metabolic pathways

RefSeq annotations of the genomes were downloaded from NCBI (Sayers et al., 2024) and the proteomes of the strains uploaded to the BlastKOALA tool (KEGG release v. 104.0, accessed March 2026) (Kanehisa et al., 2016), which assigns KEGG Orthology (KO) identifiers based on sequence similarity to a curated set of reference genomes (Kanehisa & Goto, 2000). The resulting KO numbers were uploaded to the KEGG Mapper v5 - Reconstruct tool (based on KEGG database v. 116.0, accessed March 2026) (Kanehisa et al., 2022) for reconstructing the sulfur (map00920) and cysteine/methionine (map00270) metabolic pathways.

### 2.5 Statistical methods

We used general linear model univariate analysis of variance (ANOVA; in SPSS Statistics 29.0.2.0, IBM) to assess differences in the total and individual VSC emissions (integrated over the whole culturing period) between different bacterial strains and the coculture. We used Tukey’s honestly significant difference (HSD) post hoc test for total and individual VSC emissions, followed by a Bonferroni correction to account for multiple testing. All data were log_10_-transformed to approach normal distribution.

## 3. Results

### 3.1 Differences in VSC production by bacterial cultures

*Oceanobacillus* sp. was the most prolific producer of VSCs followed by *N. aurantiaca*, producing circa 1000 and 300 pmol mL^-1^, respectively, in total during the culturing period (**Fig. 1a**). *A. arnarulunnguaqae* produced orders of magnitude less VSCs compared to the other two strains, and only 70% of the amount produced by the coculture (**Fig. 1a**). It should be noted that *Oceanobacillus* sp. did not grow as well as the other two strains (**Fig. S1a**), so the growth-normalized emission rates distinguish *Oceanobacillus* sp. from the others even further (**Fig. S1b**). Coculture emissions were a combination of the main component *A. arnarulunnguaqae* (95% of the viable cells at the end of the experiment) and *N. aurantiaca* (4% of the viable cells).

**Fig. 1.**
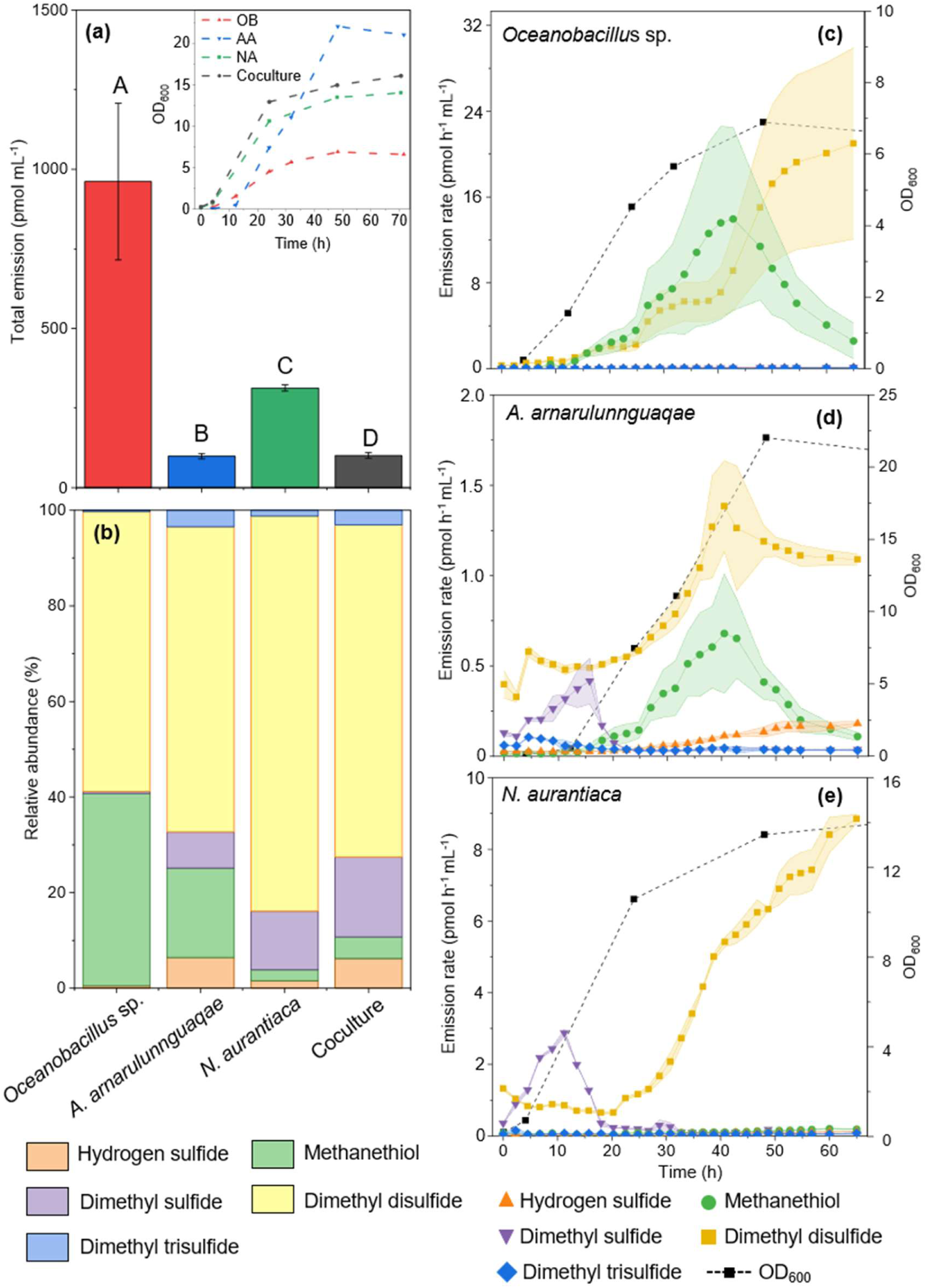
Total and real-time production of VSCs by the three bacterial strains during the 68h culturing period. **(a)** Total VSC emissions (mean ± SE, N=3) integrated over the culture period; **(b)** Relative abundances (%) of individual VSCs out of the total emission; Production profiles (mean ± SE, N=3) and the corresponding growth curves for **(c)** *Oceanobacillus* sp.; **(d)** *A. arnarulunnguaqae*; **(e)** *N. aurantiaca*. Insert in **(a)** shows the growth curves of each strain, and capital letters indicate the Tukey’s HSD ANOVA post hoc results between different strains, with differing letters meaning significant differences (*p* < 0.05)

DMDS was the most abundantly produced compound in all cultures (**Fig. 1b**), but for *Oceanobacillus* sp. more than the other strains (**Fig. S2a**), circa 600 pmol mL^-1^ in total during the culture period. *Oceanobacillus* sp. also produced orders of magnitude more MeSH than the others (**Fig. S2a**), circa 400 pmol mL^-1^. In contrast, *A. arnarulunnguaqae* and *N. aurantiaca* produced relatively more H_2_S, DMS, and DMTS (**Fig. 1b**), although the absolute concentrations were similar between all strains for H_2_S and DMTS (**Fig. S2b**). *N. aurantiaca* produced higher concentrations of DMS than the other strains (**Fig. S2b**), circa 40 pmol mL^-1^. *N. aurantiaca* also produced DMDS more abundantly than *A. arnarulunnguaqae* and the coculture (**Fig. S2a**), circa 250 pmol mL^-1^. VSC composition of the coculture was a combination of all the compounds produced by the different strains individually (**Fig. 1b** and **Fig. S2ab**).

### 3.2 Sulfur metabolic pathways and real-time production of VSCs during growth

*Oceanobacillus* sp., *A. arnarulunnguaqae*, and *N. aurantiaca* genomes matched total of 17, 16, and 8 orthologs in the sulfur metabolism pathways, respectively, and 34, 23, and 21 orthologs in the cysteine and methionine metabolism pathways, respectively (**Table S5**). In general, VSCs showed similar trends between the three strains (**Fig. 2**), where (1) H_2_S increased steadily throughout the culturing, (2) DMS peaked at lag-phase, (3) MeSH peaked in late active phase but rapidly decreased afterwards, (4) DMDS also peaked at late active phase but decreased slowly or increased in stationary phase, and (5) DMTS was produced at constant, very low rates throughout the culturing with no obvious trends relating to bacterial growth phases.

**Fig. 2.**
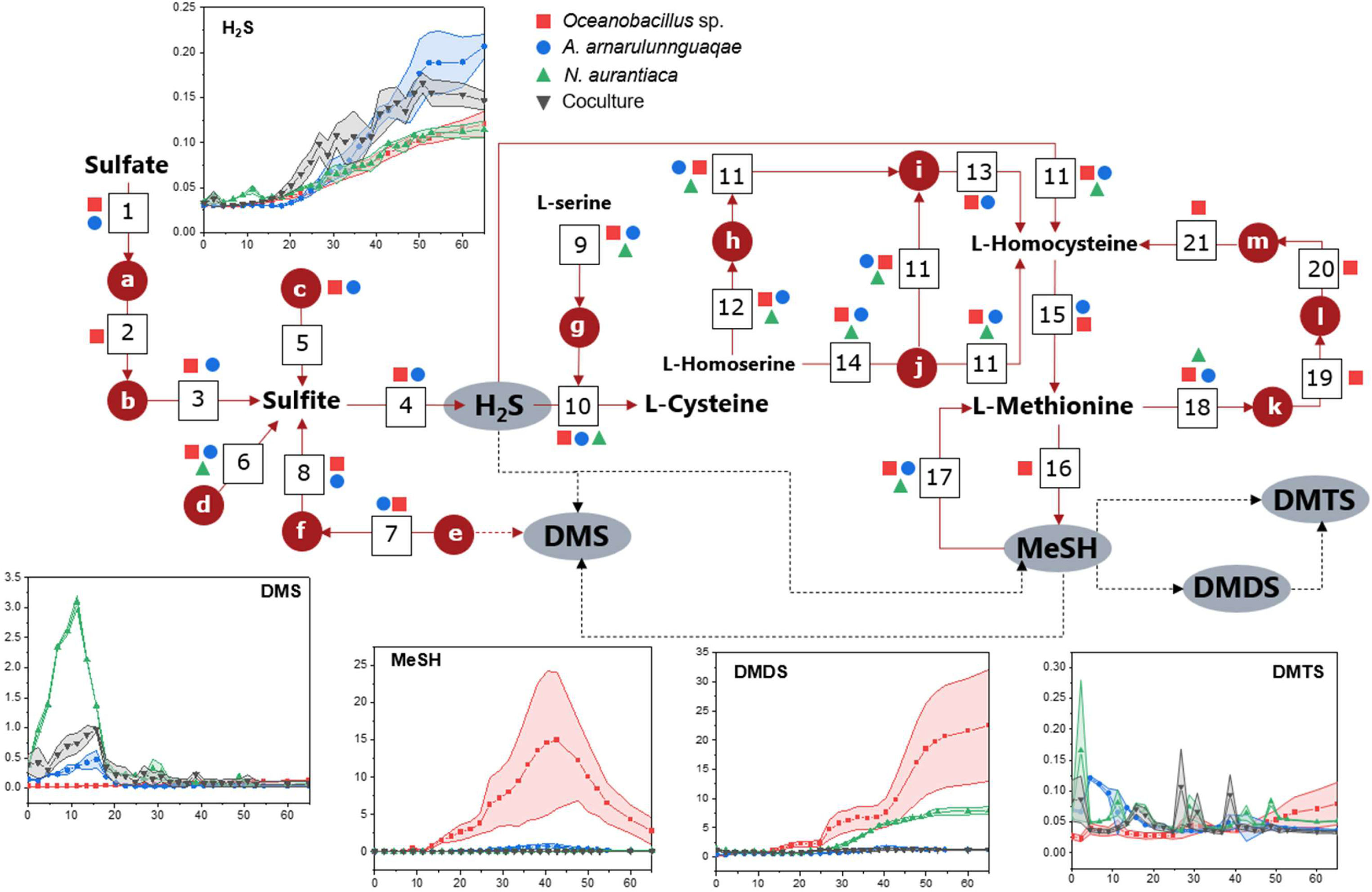
Sulfur metabolic pathways of *Ocenobacillus* sp., *A. arnarulunnguaqae*, and *N. aurantiaca* involved in VSC production. Grey ellipses indicate VSCs and bold-letter compounds major metabolic products. Red circles with letters represent minor intermediates, and squares with numbers represent enzymes. Continuous arrows indicate pathways with confirmed intermediates and enzymes, and dashed arrows possible connections, e.g., through methylation/oxidation. Symbols next to enzymes indicate they were identified from the corresponding strain. X-axis represent time (h) and y-axis emission rates (pmol h^-1^ mL^-1^). The pathway information was obtained from the KEGG database (Kanehisa & Goto, 2000). Names of enzymes and intermediates are shown in **Table 1**.

**Table 1.** Definitions for numbers and letters in Fig.2. Enzyme commission (EC) numbers indicate the corresponding enzyme-catalyzed reaction.

| Number | Enzyme | Letter | Intermediate |
| --- | --- | --- | --- |
| 1 | Sulfate adenylyltransferase (EC 2.7.7.4) | a | Adenylyl sulfate |
| 2 | Adenylyl-sulfate kinase (EC 2.7.1.25) | b | 3'-Phosphoadenylyl sulfate |
| 3 | Phosphoadenylyl-sulfate reductase (thioredoxin) (EC 1.8.4.8) | c | Alkanesulfonate |
| 4 | Assimilatory sulfite reductase (NADPH/ferredoxin) (EC 1.8.1.2/EC 1.8.7.1) | d | Thiosulfate |
| 5 | Alkanesulfonate monooxygenase (EC 1.14.14.5) | e | Dimethylsulfonate |
| 6 | Thiosulfate sulfurtransferase (EC 2.8.1.1) | f | Methanesulfonate |
| 7 | Dimethylsulfone monooxygenase (EC 1.14.14.35) | g | O-Acetyl-L-serine |
| 8 | Methanesulfonate monooxygenase (EC 1.14.14.34) | h | O-succinyl-L-Homoserine |
| 9 | Serine O-acetyltransferase (EC 2.3.1.30) | i | L-Cystathionine |
| 10 | Cysteine synthase (EC 2.5.1.47) | j | O-acetyl-L-homoserine |
| 11 | Cystathionine gamma-synthase (EC 2.5.1.48) | k | S-Adenosyl-L-methionine |
| 12 | Homoserine O-succinyltransferase (EC 2.3.1.46) | l | S-Adenosyl-L-homocysteine |
| 13 | Cysteine-S-conjugate beta-lyase (EC 4.4.1.13) | m | S-D-Ribosyl-L-homocysteine |
| 14 | Homoserine O-acetyltransferase (EC 2.3.1.31) |  |  |
| 15 | Methionine synthase (EC 2.1.1.13/EC 2.1.1.14) |  |  |
| 16 | Methionine gamma-lyase (EC 4.4.1.11) |  |  |
| 17 | O-acetylhomoserine aminocarboxypropyltransferase (EC 2.5.1.49) |  |  |
| 18 | Methionine adenosyltransferase (EC 2.5.1.6) |  |  |
| 19 | DNA (cytosine-5-)-methyltransferase (EC 2.1.1.37) |  |  |
| 20 | Adenosylhomocysteine nucleosidase (EC 3.2.2.9) |  |  |
| 21 | S-ribosylhomocysteine lyase (EC 4.4.1.21) |  |  |

For *Oceanobacillus* sp., we identified a complete assimilatory sulfate reduction pathway (KEGG module M00176), while for *A. arnarulunnguaqae* the pathway was mostly complete, missing only adenylyl-sulfate kinase (K00860, **Fig. 2**; 2). By contrast, *N. aurantiaca* had no orthologues in this pathway. The final product of this pathway is H_2_S (**Fig. 2**; sulfate → sulfite → H_2_S). For all strains, H_2_S emission rates were small (<0.2 pmol h^-1^ mL^-1^) but increased linearly with cell density from active phase onwards (**Fig. 1c-d** and **Fig. 2**). All strains also had a complete cysteine biosynthesis pathway from serine (M00021).

For *Oceanobacillus* sp., we identified complete methionine biosynthesis (M00017) and degradation (M00035) pathways. Its genome also encoded for mostly complete pathways for cysteine biosynthesis from methionine (M00609; 4/6 orthologs present) and for methionine salvage (M00034; 5/8 orthologs present). The genome contained an ortholog of methionine gamma-lyase (K01761), which cleaves MeSH from methionine (**Fig. 2**; 16). *A. arnarulunnguaqae* had a complete and mostly complete methionine degradation (M00035) and synthesis (M00017; 6/7 orthologs present) pathways, respectively. Both were incomplete in *N. aurantiaca* (3/4 and 5/7 orthologs present, respectively). The methionine salvage pathway was incomplete for both strains (M00034, 2/8 orthologs present). For these two strains, methionine adenosyltransferase (K00789) instead of methionine gamma-lyase (K01761) was identified in the methionine degradation pathway (**Fig. 2**; 18), which does not lead directly to the production of MeSH. This enzyme was also identified for *Oceanobacillus* sp. For all three strains, we also identified an O-acetylhomoserine aminocarboxypropyltransferase (K01740), which catalyzes the reaction between O-acetyl-L-homoserine and MeSH (**Fig. 2**; 17), potentially leading to consumption of MeSH. MeSH can also be produced via methylation of H_2_S (**Fig. 2**: H_2_S → MeSH, dashed lines).

MeSH production by *Oceanobacillus* sp. increased in the exponential phase to 15 pmol h^-1^ mL^-1^ but rapidly decreased when nearing stationary phase (**Fig. 1c**). MeSH production by *A. arnarulunnguaqae* was much smaller (max. 0.8 pmol h^-1^ mL^-1^) but had the same pattern (**Fig. 1d**). *N. aurantiaca* produced MeSH at very low levels (<0.3 pmol h^-1^ mL^-1^) and had no obvious pattern (**Fig. 1e**), similar to the coculture (**Fig. S3).**

DMS production by *A. arnarulunnguaqae* and *N. aurantiaca* peaked early in the exponential phase (max. 0.5 and 3 pmol h^-1^ mL^-1^, respectively) and decreased rapidly thereafter (**Fig. 1d-e**). Similar trend was observed for the coculture (**Fig. S3**). *Oceanobacillus* sp. produced only small amounts (<0.2 pmol h^-1^ mL^-1^) of DMS with an increasing trend towards the stationary phase (**Fig. 1c** and **Fig. 2**). We did not identify any pathways leading to the production of DMS for any of the strains in the KEGG analysis, but DMS can be produced from methylation of MeSH or H_2_S (**Fig. 2**; MeSH → DMS and H_2_S → DMS, dashed lines).

DMDS production by *Oceanobacillus* sp. increased significantly in the exponential phase and continued to the stationary phase (max. 23 pmol h^-1^ mL^-1^; **Fig. 1c**). Same trend was observed for *N. aurantiaca* (**Fig. 1e**), although in smaller amounts (max. 9.5 pmol h^-1^ mL^-1^). For *A. arnarulunnguaqae*, DMDS decreased after peaking at the end of the exponential phase (max. 1.6 pmol h^-1^ mL^-^1; **Fig. 1d**). Coculture had a slight increasing trend towards the stationary phase (**Fig. S3**). DMDS is not included in the KEGG sulfur metabolic maps but can be biologically and abiotically oxidized from MeSH (**Fig. 2**; MeSH → DMDS, dashed line).

### 3.3 Emission of VSCs from thawing permafrost

Permafrost soil samples emitted small amounts of VSCs during the whole thaw period (2.3 ± 0.06 pmol g^-1^; **Fig. 3a**), with DMS and DMDS as the largest contributors (37 and 30%, respectively), and MeSH and DMTS present in smaller amounts (23 and 10%, respectively; **Fig 3b**). H_2_S was not observed above detection limit. Permafrost soil samples released VSCs from the beginning of the thaw, with steadily declining emission rates for circa 20 h, followed by a short period of low emissions (**Fig. 3c**). After this, emission rates started increasing again, with DMS, DMDS, and DMTS showing rapid increase in the latter part of the thawing period, while MeSH increased more slowly. At the end of the 40-h thawing period, VSC emissions were still increasing (**Fig. 3c**). Permafrost VSC emission profile was thus divided into three main phases: release, lag, and production (**Fig. 3d**).

**Fig. 3.**
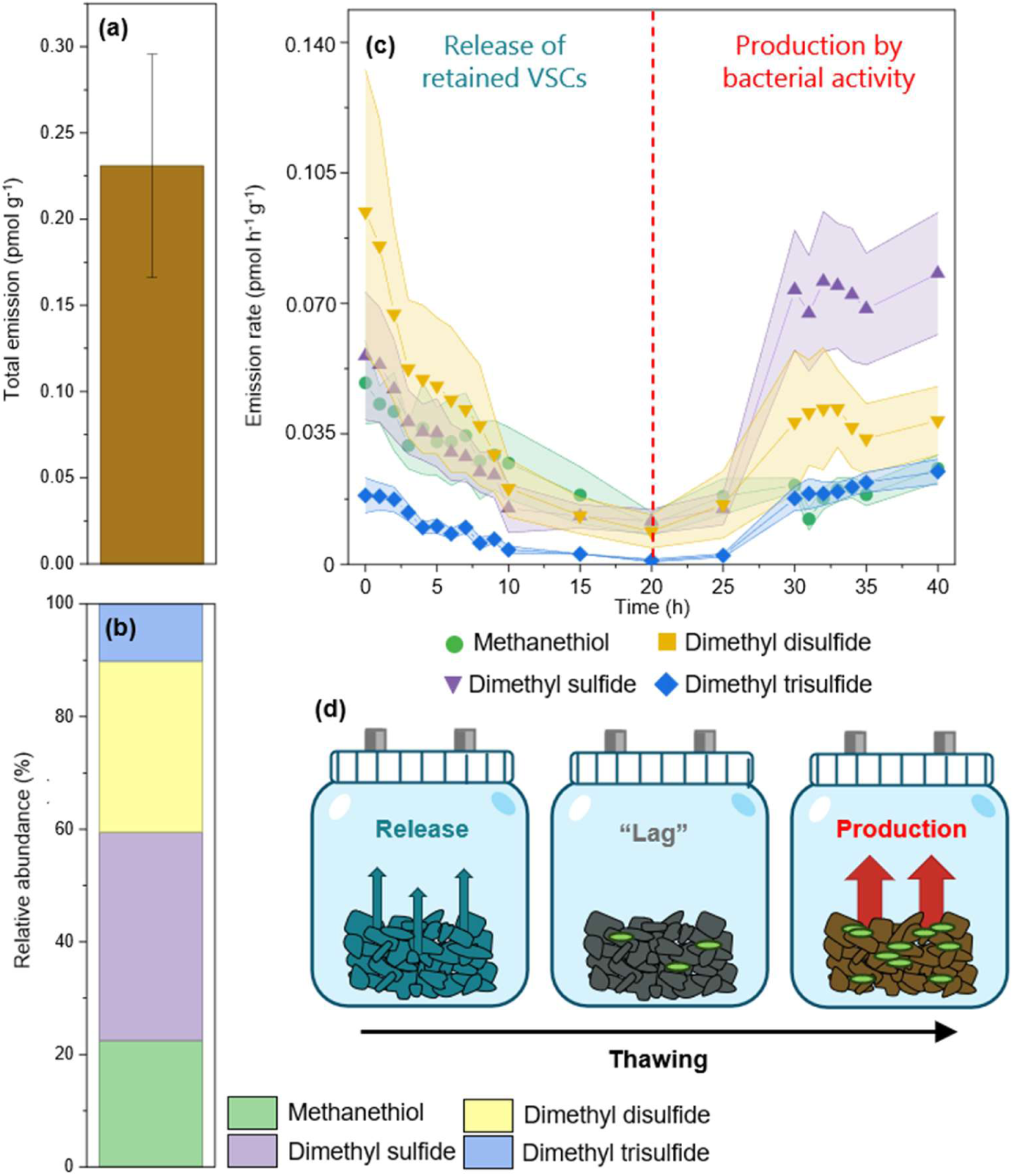
Total and real-time production of VSCs from permafrost during thawing. **(a)** Total VSC emission (mean ± SE, N=6) integrated over the thaw period; **(b)** Relative abundances (%) of individual VSCs out of the total emission; **(c)** Emission profile (mean ± SE, N=6) for thawing permafrost; **(d)** Schematic representation of possible processes responsible for the three main phases observed in the permafrost emission profile during thawing: release of VSCs retained in the permafrost matrix, lag phase of low emissions, and *de novo* production of VSCs by increased bacterial activity.

## 4. Discussion

### 4.1. VSC emissions from bacterial cultures and thawing permafrost suggest increased trends in the future

As hypothesized, we observed all three strains produce a variety of VSCs when inorganic and organic sulfur is available in the environment. Less than 0.1% by weight of the nutrient media was sulfur-containing amino acids and only 0.1% by weight MgSO_4_, yet the studied bacteria produced significant amounts of VSCs – especially *Oceanobacillus* sp. Production of VSCs from small amount of nutrients may also be the case in their natural environment, if conditions are favorable for increased microbial activity, such as when permafrost thaws.

We hypothesized that thawing permafrost emits quantifiable amounts of reduced VSCs, originating at least in part from microbial activity. Our results from permafrost incubations support this hypothesis: We show that permafrost emits VSCs during thaw, with an initial release of trapped compounds from previous microbial production within the permafrost matrix, followed by a secondary, accelerated emission phase, presumably when bacterial activity increases (**Fig. 3c-d**). In a previous study reporting non-sulfur VOC production in the same measurements, rapid emission of methanol and ethanol was observed within the first few hours of thaw, but the emissions declined thereafter without any secondary emission pattern (Kramshøj et al., 2018). The authors hypothesized that this was due to release of previously trapped gases rather than *de novo* microbial production. In another study, an initial burst of trapped CH_4_ was observed upon thaw of permafrost samples, but after 48 h, methanotrophs were shown to actively consume CH_4_ and produce CO_2_ when energy sources became available during thaw (Mackelprang et al., 2011).

While the coculture was dominated by *A. arnarulunnguaqae* at the end of the experiment (95% relative abundance), the volatile composition differed from the pure *A. arnarulunnguaqae* cultures, indicating that the other two strains – but especially *N. aurantiaca* – contributed to the volatile composition, although only present in small amounts (4% relative abundance). This demonstrates that the VSCs released from a complex mixture are not directly the sum of the mixture’s individual components, as bacterial functions may differ when species grow together, and emission patterns cannot be estimated from microbial composition alone. Our results also show that lower levels of VSCs were not caused by poor growth, as *A. arnarulunnguaqae* exhibited the highest growth rate in the chosen conditions but lowest emission rates, while *Oceanobacillus* sp. grew poorest but produced the highest levels of VSCs. This means that optimal growth is not always the driver behind VSC emissions, but rather depend on the microbial community composition and nutrient availability. For example, sulfate-reducing bacteria can contribute significantly to carbon flow, and sustain transcriptional activity at extremely low growth rates even as minor constituents of the community (Hausmann et al., 2016; Pester et al., 2010). Consequently, even in extreme environments – like permafrost – VSC production may increase if suitable nutrients become available, regardless of growth rates.

Organic matter trapped inside permafrost contains the remnants of past vegetation and animals, which in the case of Greenland were part of a rich boreal forest ecosystem 2 million years ago (Kjær et al., 2022). This organic matter also likely contains vast amounts of sulfur-containing amino acids, which become available to microbial activity due to ongoing climate change. Thawing of permafrost accelerates organic matter turnover, changes redox conditions, and enhances sulfide oxidation (Bataille & Skierszkan, 2025; Kemeny et al., 2023). Thus, more inorganic sulfur is also mobilized in the thawed soil for the use of sulfate reducing bacteria, the amounts of which are significantly increased during thawing (Li et al., 2020; Stackhouse et al., 2015). This combination of increased, available sulfur and higher activity of sulfur-utilizing bacteria is likely to result in higher emissions of VSCs and CO_2_ from permafrost-affected areas in the future, unless the microbiome can cycle these reactive gases before they diffuse to the atmosphere (Rinnan, 2024).

### 4.2. Amino acid metabolism and assimilatory sulfate reduction behind VSC emissions

MeSH was produced by all strains, but most significantly by *Oceanobacillus* sp. (**Fig. 1b**). It is a direct degradation product of L-methionine, but can also be produced from H_2_S following further methylated into DMS (Carrión et al., 2015). Complete methionine biosynthesis and mostly complete degradation and salvage pathways – including methionine gamma-lyase – were identified for *Oceanobacillus* sp. (**Fig. 2**; 16), with the continuous production profile (**Fig. 1c**) showing accelerated production of MeSH in the exponential phase and suggesting efficient use of methionine in the nutrient media. The genes encoding methionine gamma-lyase are upregulated in methionine rich conditions in cheese-ripening bacteria (Sato & Nozaki, 2009), which may also explain our observations for decreasing concentration of MeSH in stationary pahse: When methionine diminishes, growth rates slow down and methionine gamma-lyase is downregulated, stopping MeSH production. Consequently, in *Oceanobasillus* sp., MeSH is tightly linked to active growth. Same connection has been shown for human pathogenic, VSC-producing bacteria (Roslund et al., 2020). Furthermore, we showed previously that salt-stress affects a plethora of volatile metabolites (Salinas-García et al., 2025), including MeSH, which may be explained by hindered active growth.

For *A. arnarulunnguaqae* and *N. aurantiaca*, methionine gamma-lyase was not identified in the methionine degradation pathway, but instead an alternative enzyme (methionine adenosyltransferase) (**Fig.2**; 18). This enzyme – also present in *Oceanobasillus* sp. – catalyzes a reaction not directly leading to the production of MeSH. This difference is likely the reason behind the low MeSH production by *A. arnarulunnguaqae* and *N. aurantiaca* (**Fig. 1d-e**), although methionine was readily available. These differences may originate from differing adaptations to the natural environment. For *A. arnarulunnguaqae*, low temperatures and nutrient scarcity in permafrost soil may lead to sulfur being conserved rather than emitted as VSCs.

DMDS was also produced by all strains, although much less by *A. arnarulunnguaqae* compared to the other two strains. For *Oceanobacillus* sp. and *N. aurantiaca*, DMDS production accelerated in the stationary phase, suggesting a metabolic pathway expressed after active growth has slowed, unlike for MeSH, linked to the exponential phase. Same trend has been previously observed for human pathogenic, VSC-producing bacteria (Roslund et al., 2020). While not well studied, the production of DMDS by bacteria has been linked to degradation of methionine leading to MeSH, followed by biological or abiotic oxidation to DMDS, or through S-methylcysteine without the MeSH intermediate (Tomita et al., 1987). While production capacities differed between our studied strains, DMDS was exclusively connected to the later growth stages, and therefore, secondary metabolism. DMDS production can have antagonistic effects on competing organisms, such as yeasts (Rahmath et al., 2025), so in soil it may be advantageous for the bacteria, while in environments like permafrost, where fungi are rare, there is less need for competition (Gittel et al., 2013). This may explain why the permafrost strain, *A. arnarulunnguaqae,* produced significantly less DMDS compared to the two strains isolated from the active layer.

DMS was produced by all strains but most abundantly by *N. aurantiaca.* Unlike the other VSCs discussed here, DMS cannot be directly produced from amino acids. Methionine or cysteine must be first catabolized to either H_2_S or MeSH, and these compounds further methylated to DMS. Complete methionine degradation pathways were found for all strains, although the enzymes related to them differed, as discussed previously. The enzymes related to methylation of H_2_S and MeSH – MddA and MddH – are recognized in both aquatic and terrestrial environments as significant alternatives for the DMSP/MMPA pathway (Carrión et al., 2015; Zhang et al., 2024). While these enzymes were not identified for the studied strains, when working with novel bacteria, some orthologs may be missed if their sequences are not similar enough to the curated database. In addition, the genomes used in this study were on the contig level and some genes may have been cut, preventing recognition in BlastKOALA. Consequently, some pathways may appear incomplete, missing one or two enzymes, but actually present in the bacterial genome. In our study, the nutrient media did not contain DMSP, and therefore, DMS production by is likely unrelated to it. Additionally, sulfate reduction pathways were not identified for *N. aurantiaca* – the main producer – so DMS is unlikely to originate from inorganic sulfur either. Amino acid degradation and subsequent methylation are, therefore, the most likely precursors for DMS in these soil and permafrost bacteria, but further genome analysis together with isotope tracing should be done in the future to confirm this.

Production of DMS differed from the other VSCs as it spiked in the early exponential phase and stopped when the production of other VSCs started. This suggests that DMS is not necessarily linked to active growth, but to a pathway only expressed as growth starts. The rapid diminishing may be due to bacteria switching to (1) metabolic routes related to active growth that do not produce DMS, or (2) oxidization of DMS to DMSO, which has been documented for marine bacteria (Green et al., 2011; Lidbury et al., 2016), but not in this study. The latter may be used for detoxification or converting DMS to a more accessible – less volatile and more soluble – form.

Assimilatory sulfate reduction pathway was identified for *Oceanobacillus* sp. and *A. arnarulunnguaqae*, but none of the strains had genes for dissimilatory sulfate reduction. Assimilatory sulfate reduction incorporates sulfur into organic molecules to produce sulfur-containing amino acids, and therefore, the pathway is not thought to release H_2_S, unlike the dissimilatory pathway where H_2_S is a main product (Jia et al., 2026). Our results suggest that the studied bacteria use inorganic sulfur to produce amino acids, rather than energy, and the observed H_2_S emissions do not originate directly from sulfate. Instead, H_2_S can be produced from the cleavage of L-cysteine, which was available in the nutrient media. Sulfate reduction pathway was not identified for *N. aurantiaca* but it still produced H_2_S, which further supports cysteine as the main precursor rather than sulfate. While H_2_S itself can be a precursor for the formation of other VSCs in bacterial metabolism, this does not seem to be the case for the studied strains because the production of H_2_S increased throughout the culturing, independent of MeSH and DMS. This further suggests that H_2_S has different metabolic precursors (L-cysteine or sulfate) compared to the other VSCs (L-methionine and S-methylcysteine), and the production likely continues throughout the bacterial growth because these nutrients are not as quickly depleted.

Overall, our findings suggest that Arctic soil bacteria are abundant producers of VSCs when sulfur is readily available, and that thawing increases VSC emissions from permafrost soil, suggesting higher input of sulfur into the atmosphere from the warming Arctic in the future. Our results encourage future efforts in assessing the importance of soil microbial VSCs in the Arctic – including but not limited to DMS – utilizing, e.g., mesocosm or artificial soil community approaches.

## Acknowledgement

The Danish National Research Foundation (Center for Volatile Interactions - VOLT, DNRF168), the European Research Council (ERC) under the European Union’s Horizon 2020 research and innovation programme (grant agreement no. 771012), and Jenny and Antti Wihuri Foundation financed this work. We would also like to thank Bo Elberling for collecting the permafrost samples and Magnus Kramshøj and Thomas Holst for conducting the permafrost measurements.

## Data availability

Upon acceptance, full data is available at the Electronic Research Data Archive at the University of Copenhagen (ERDA, UCPH) through a permanent DOI link.

## Author contribution

KR, MSG, AP, and RR conceived the idea and designed the study; KR and MSG planned and conducted the measurements; KR processed and analyzed the VSC data and conducted statistical analysis; KR and MSG conducted pathway analysis; RR and AP provided supervision; KR led the writing of the manuscript. All authors reviewed and contributed critically to the drafts and gave final approval for publication.

---

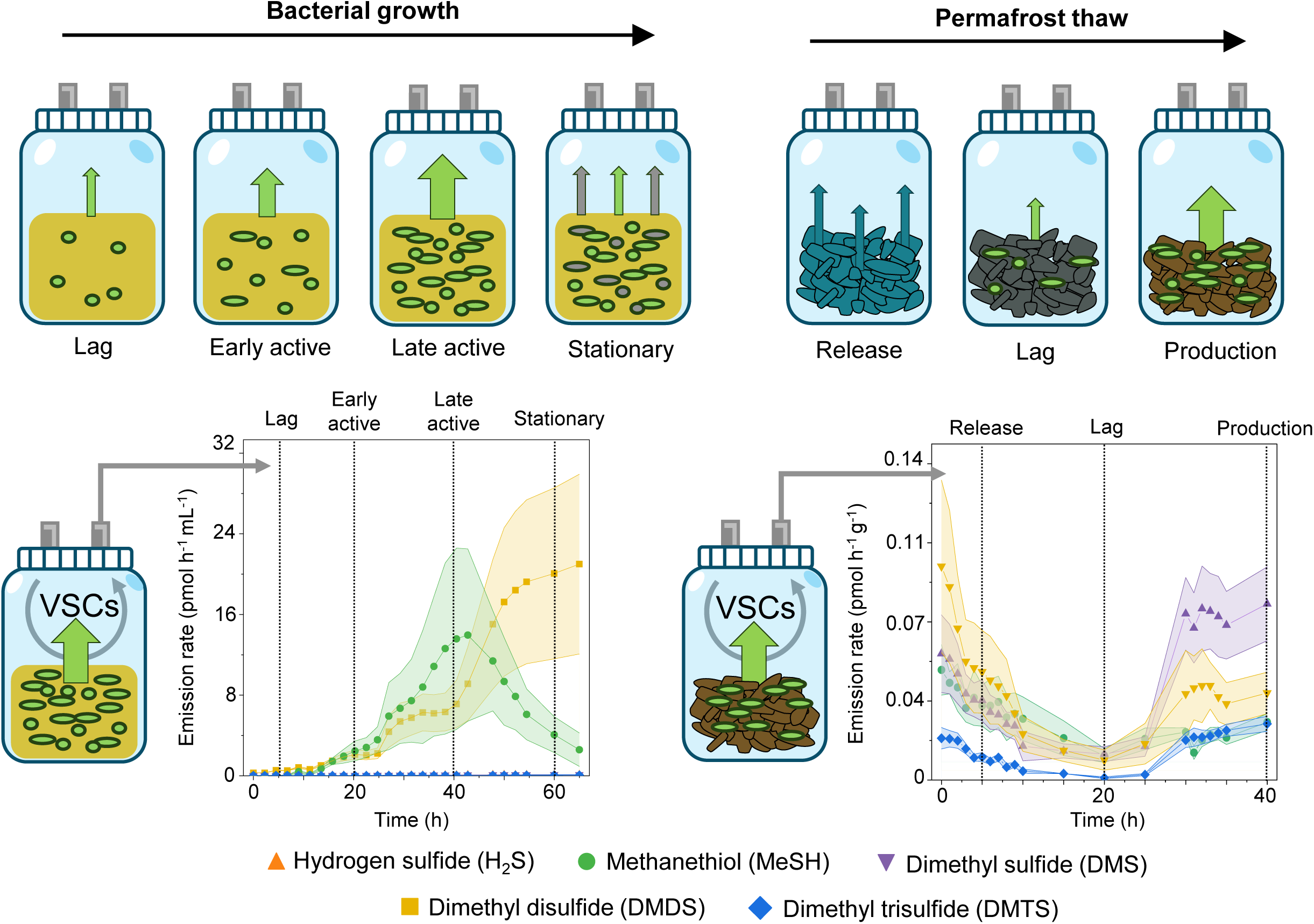

